# Comparative transcriptome analyses reveal genes associated with SARS-CoV-2 infection of human lung epithelial cells

**DOI:** 10.1101/2020.06.24.169268

**Authors:** Darshan S. Chandrashekar, Upender Manne, Sooryanarayana Varambally

**Author notes:** Correspondence to: Sooryanarayana Varambally, Ph.D., Molecular and Cellular Pathology, Department of Pathology, Wallace Tumor Institute, 4^th^ floor, 20B, University of Alabama at Birmingham, Birmingham, AL 35233, USA, And Darshan S. Chandrashekar Ph.D., Department of Pathology, University of Alabama at Birmingham, Birmingham, AL. Share Senior Authorship (UM).

## Abstract

Understanding the molecular mechanism of SARS-CoV-2 infection (the cause of COVID-19) is a scientific priority for 2020. Various research groups are working toward development of vaccines and drugs, and many have published genomic and transcriptomic data related to this viral infection. The power inherent in publicly available data can be demonstrated via comparative transcriptome analyses. In the current study, we collected high-throughput gene expression data related to human lung epithelial cells infected with SARS-CoV-2 or other respiratory viruses (SARS, H1N1, rhinovirus, avian influenza, and Dhori) and compared the effect of these viruses on the human transcriptome. The analyses identified fifteen genes specifically expressed in cells transfected with SARS-CoV-2; these included *CSF2* (colony-stimulating factor 2) and *S100A8* and *S100A9* (calcium-binding proteins), all of which are involved in lung/respiratory disorders. The analyses showed that genes involved in the Type1 interferon signaling pathway and the apoptosis process are commonly altered by infection of SARS-CoV-2 and influenza viruses. Furthermore, results of protein-protein interaction analyses were consistent with a functional role of CSF2 in COVID-19 disease. In conclusion, our analysis has revealed cellular genes associated with SARS-CoV-2 infection of the human lung epithelium; these are potential therapeutic targets.

## Introduction

Infection of severe acute respiratory syndrome coronavirus-2 (SARS-CoV-2) is the cause of human coronavirus disease 2019 (COVID-19). The recent pandemic has caused devastation due to rapid spread of this viral infection. As a respiratory illness, the disease is readily transmitted. It also has a long incubation and can be carried asymptomatically, thus spreading through communities (1). The COVID-19 pandemic has affected almost every country regardless of their medical infrastructure and economic status. It has caused a healthcare crisis and created a devastating economic burden, including high unemployment, which has exacerbating the effect of the disease (2). At present, more than 9.4 million people from 213 countries have been infected with the virus [https://www.worldometers.info/coronavirus/]. The rapid spreading of this respiratory infection has forced millions to shelter in their homes and has led to death of more than 480,000 individuals. Additionally, COVID-19 disproportionately affects patients, particularly minorities in the U.S. and those with chronic problems such as hypertension, lung disease, diabetes, and immunocompromised conditions.

In the last three decades, the world has witnessed zoonotic transmission of various viruses (from animals to humans) leading to severe respiratory complications. These include H1N1, avian influenza, severe acute respiratory syndrome (SARS), and Middle East respiratory syndrome coronavirus (MERS□CoV) (3, 4). Although infections of these viruses is often fatal, their effect is restricted to geographic locations such as Africa, Asia, and South America. As the world awaits a vaccine for SARS-CoV-2, efforts are being made to understand the molecular mechanisms of these infections (5, 6).

High-throughput technologies such as RNA sequencing and microarrays are useful in the detection of respiratory virus infections and in understanding their molecular effect on human lung epithelial cells (7). Extensive data on corona virus sequencing has been deposited in public repositories such as NCBI Gene Expression Omnibus (GEO) and EMBL Array Express (8, 9). Meta-analysis and mining of such data can aid in a) understanding the molecular impact of COVID-19, b) elucidating differences and similarities between SARS-CoV-2 and other respiratory virus infections, and c) identification of targets for drug development. In the current study, we performed comparative analysis of publicly available gene expression data related to human lung epithelial cells infected with a respiratory virus. The analyses identified genes specifically expressed by SARS-CoV-2 infection and those that are commonly altered due to infection of coronovirus-2 and/or other respiratory viruses. In particular, expression of *CSF2* (colony-stimulating factor 2) appears to be involved in COVID-19 disease.

## Methods

### RNA sequencing data analysis

A search of the NCBI GEO database for microarray or RNA-sequencing data publicly available as of 13^th^ April 2020, found RNA-seq data uploaded by Blanco-Melo et al. (GSE147507) (6). We focused on samples related to normal human bronchial epithelial cells subjected to mock treatment (n=3) or SARS-CoV-2 infection (n=3). Raw sequencing data related to selected samples of GSE147507 as fastq files were downloaded from Sequence Read Archive (SRA) using fastq-dump of sratoolkit v2.9.6 [http://ncbi.github.io/sra-tools/]. First, raw sequencing reads were trimmed to remove adapter sequences and low-quality regions using Trim Galore! (v0.4.1) [http://www.bioinformatics.babraham.ac.uk/projects/trim_galore/]. Trimmed reads were subjected to quality control analysis using FastQC [https://www.bioinformatics.babraham.ac.uk/projects/fastqc/]. Tophat v2.1 was used to map trimmed raw reads to the human reference genome (hg38) (10). All bam files from multiple runs related to the same samples were merged and sorted using SAMtools (Version: 1.3.1) (11). Finally, raw read counts were enumerated for each gene in each sample using HTSeq-count (12).

Analysis of differential expression was performed using DESeq2 according to a standard protocol [https://bioconductor.org/packages/release/bioc/vignettes/DESeq2/inst/doc/DESeq2.html] (13). Genes with adj.P-value <0.05 and absolute fold change >= 1.5 were considered as significantly differentially expressed. Gene ontology enrichment analyses of the Differentially Expressed Genes (DEGs) were accomplished by use of the Database for Annotation, Visualization and Integrated Discovery (DAVID) v6.8 online tool (14). Gene ontology (GO) biological processes with P-values <0.05 and gene counts >2 were considered as significantly enriched.

### Microarray data collection and analysis

The NCBI GEO database was queried for microarray data related to SARS-CoV infections of human lung epithelial cells. A query *(SARS-CoV) AND “Homo sapiens” [porgn] AND (“gse” [Filter] AND (“Expression profiling by array” [Filter]))* led to 15 search results. After screening, two studies (GSE47962, GSE17400) were selected. To find microarray data related to SARS-CoV infections of human lung epithelial cells, the GEO database was queried using *(((Human lung epithelium) OR (Human bronchial epithelial) AND “Homo sapiens” [porgn] AND (“gse” [Filter] AND “Expression profiling by array” [Filter]))) AND (viral infection AND (“gse” [Filter] AND “Expression profiling by array” [Filter])) AND (“gse” [Filter] AND “Expression profiling by array” [Filter]) AND (“Expression profiling by array” [Filter])*. This led to 38 search results, three of which (GSE49840, GSE71766, and GSE48575) were selected for analysis. **Table 1** provides sample, platform, and cell line details for all five studies.

**Table 1:**
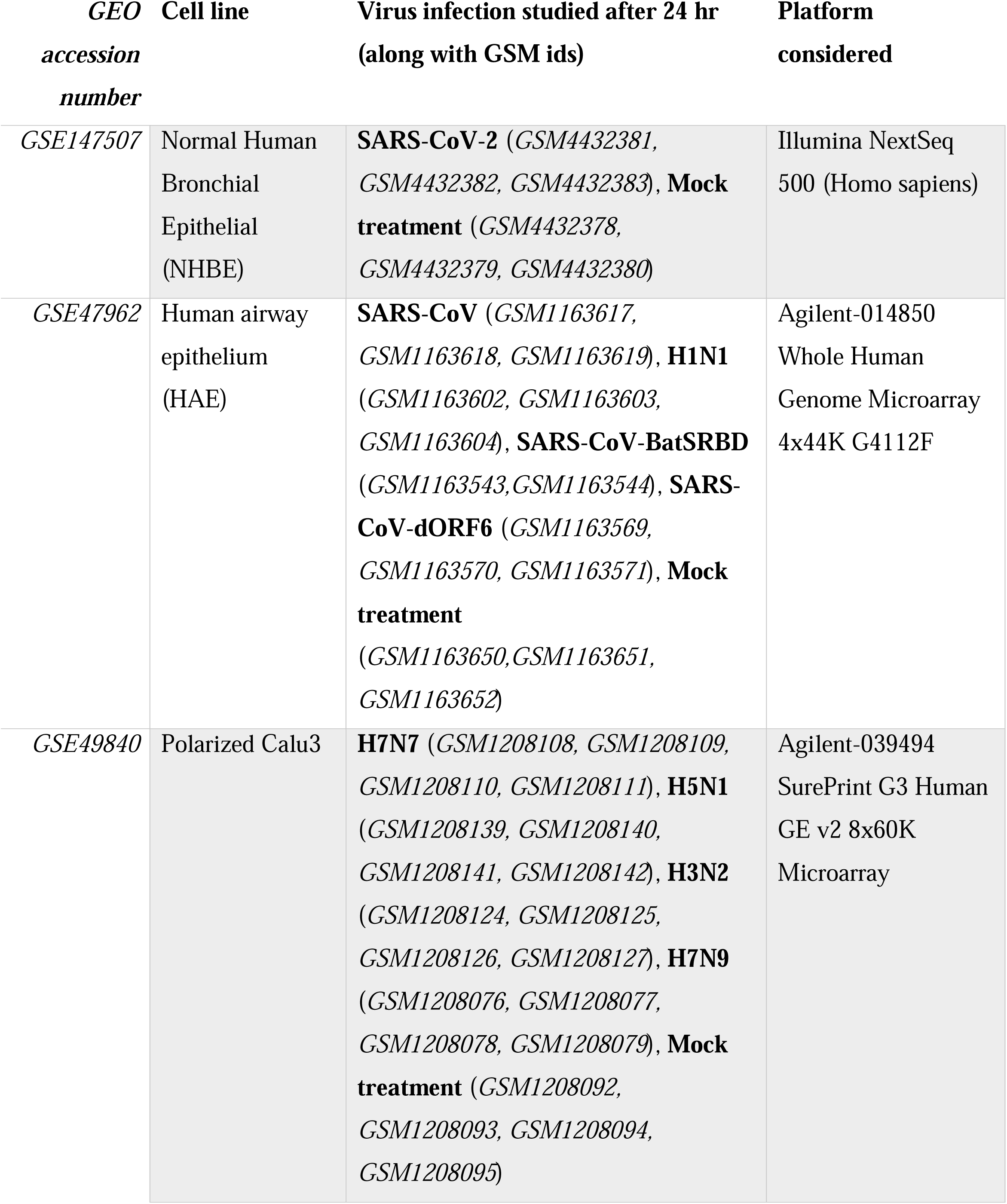
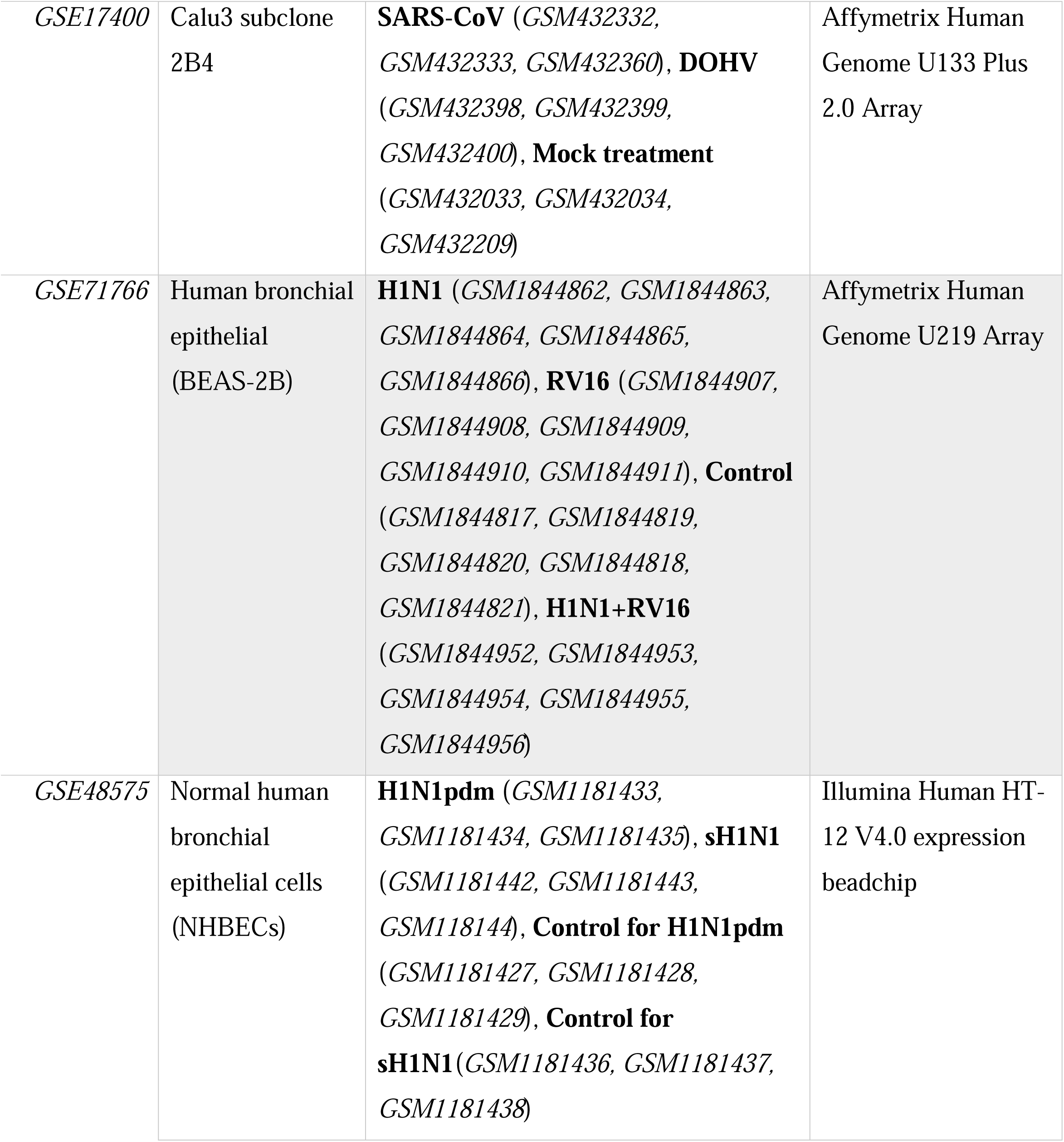
Summary of publicly available, high-throughput experimental data considered in the current study.

Since SARS-CoV-2 RNA-seq data included transcriptome profiling after 24 hours of infection, in all microarray studies we only considered samples after 24 hours of viral infection. GSE47962 included samples from human airway epithelium (HAE) cells infected with SARS-CoV, influenza virus (H1N1), or variants of SARS-CoV (SARS-dORF6 and SARS-BatSRBD) (15). GSE71766 comprised human bronchial epithelial cells (BEAS-2B) infected with rhino virus (RV), influenza virus (H1N1), or both (RV + H1N1) (16). Bronchial epithelial cell line 2B4 (a clonal derivative of Calu-3 cells) infected with SARS-CoV or Dhori virus (DOHV) were part of GSE17400 (17). GSE49840 included polarized calu-3 (cultured human airway epithelial cells) infected with human influenza virus (H3N2) or avian influenza viruses (H7N9, H5N1, and H7N7) (18). GSE48575 consisted of normal human bronchial epithelial cells (NHBEC) infected with seasonal H1N1 influenza A (sH1N1) or pandemic H1N1 influenza A (H1N1pdm) (19). GEO2R was used to identify differentially expressed genes for each of these studies independently (9). Probes with adj. P-value <0.05 and absolute fold change >=1.5 were considered as statistically significant and compared with DEGs of SARS-CoV-2 infection from RNA-seq data.

### Protein-protein interaction analysis

STRING, a database of known or predicted protein-protein interactions (PPIs) was used to obtain interactions between genes altered on SARS-CoV-2 infection (20). Output from the STRING database was uploaded to Cytoscape v3.7.2 in simple interaction format, and the Cytohubba app was employed to identify hub genes (21-23). The top 50 genes were obtained separately from the PPI network based on three network parameters (closeness, degree, and betweenness), then common genes among these were selected as hub genes. We also checked the DrugBank database to determine if a drug is available to target them (24).

## Results

### RNA sequencing identified genes altered on SARS-CoV-2 infection of normal human bronchial epithelial cells (NHBEC)

Differential expression analysis was performed for NHBEC subjected to SARS-CoV-2 infection (n=3) or mock infection (n=3) from GSE147507. In total, 164 genes were up-regulated, and 76 genes were down-regulated by SARS-CoV-2 infection compared to mock infection [**Figure 1A**]. Gene Ontology enrichment analysis of differentially expressed genes showed “Type I interferon signaling pathway”, “Inflammatory response”, “Immune response”, “Response to virus”, and “Defense response to virus” as the top 5 enriched biological processes [**Figure 1B**].

**Figure 1:**
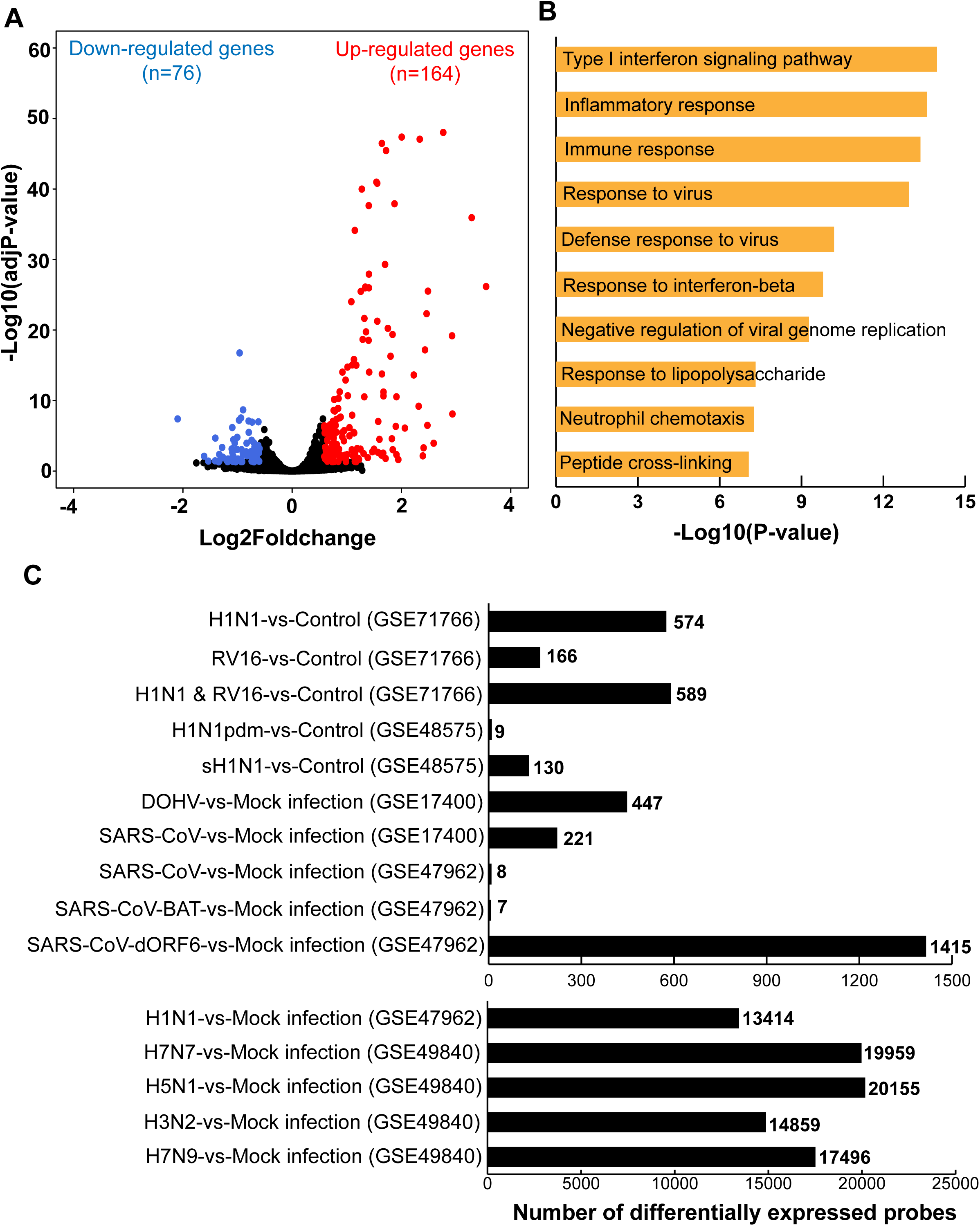
High throughput gene expression analysis of COVID-19 and other respiratory viral infection in human lung epithelial cells. (A) Volcano plot showing the proportion of differentially expressed genes after processing COVID-19 RNA-seq data [GSE147507]. (B) Top 10 biological processes associated with genes differentially expressed on COVID-19 infection of human lung epithelial cells. (C) Histogram showing the differentially expressed genes as determined by GEO2R analysis of respiratory viral infection microarray data.

### The impact of other viral infections on the human lung epithelial cell transcriptome was explored using publicly available microarray data

A search of microarray datasets in the NCBI GEO database led to identification of five studies involving human lung epithelial cells subjected to viral infection. GEO2R analyses of each study were performed separately identifying differentially expressed genes. **Figure 1C** shows DEGs identified from each comparative analysis. From GSE71766, there were 574 differentially expressed probes in H1N1-infected BEAS-2B cells compared to control; 166 probes were up-regulated by RV16 infection. The combined infection of RV16 and H1N1 altered expression of 589 probes. In the case of GSE49840, there were 19959 probes differentially expressed in H7N7-infected Calu-3 cells compared to mock infected cells; 20155, 14859, and 17496 probes were differentially expressed by H5N1, H3N2, or H7N9 infections, respectively. For GSE17400, DOHV or SARS-CoV infection of Calu-3 cells led to altered expression of 447 and 221 probes, respectively. Analysis of GSE48575 led to identification of 130 and 9 differentially expressed probes on seasonal (sH1N1) or pandemic (H1N1pdm) influenza virus infection, respectively, of NHBECs. Lastly, the processing of GSE47962 resulted in discovery of 13414, 1415, 7, and 8 differentially expressed probes after H1N1, SARS-CoV-dORF6, SARS-CoV-BAT, or SARS-CoV viral infections, respectively, of HAE cells.

### Comparative analysis of DEGs resulted in identification of SARS-CoV-2 infection-specific genes and those commonly affected by most of the viral infections

Differentially expressed genes from SARS-CoV-2 RNA-seq data were compared with GEO2R analysis results of GSE47962, GSE17400, GSE48575, GSE49840, and GSE71766. There were 15 genes differentially expressed after infection of SARS-CoV-2 [**Figure 2A**]. *TCIM, HEPHL1, TRIML2, S100A9, S100A8, CRCT1, MAB21L4, MRGPRX3*, and *CSF2 were up-regulated; HNRNPUL2−BSCL2, CXCL14, THBD, GCNT4, PCDH7*, and *PADI3* were down-regulated.

**Figure 2:**
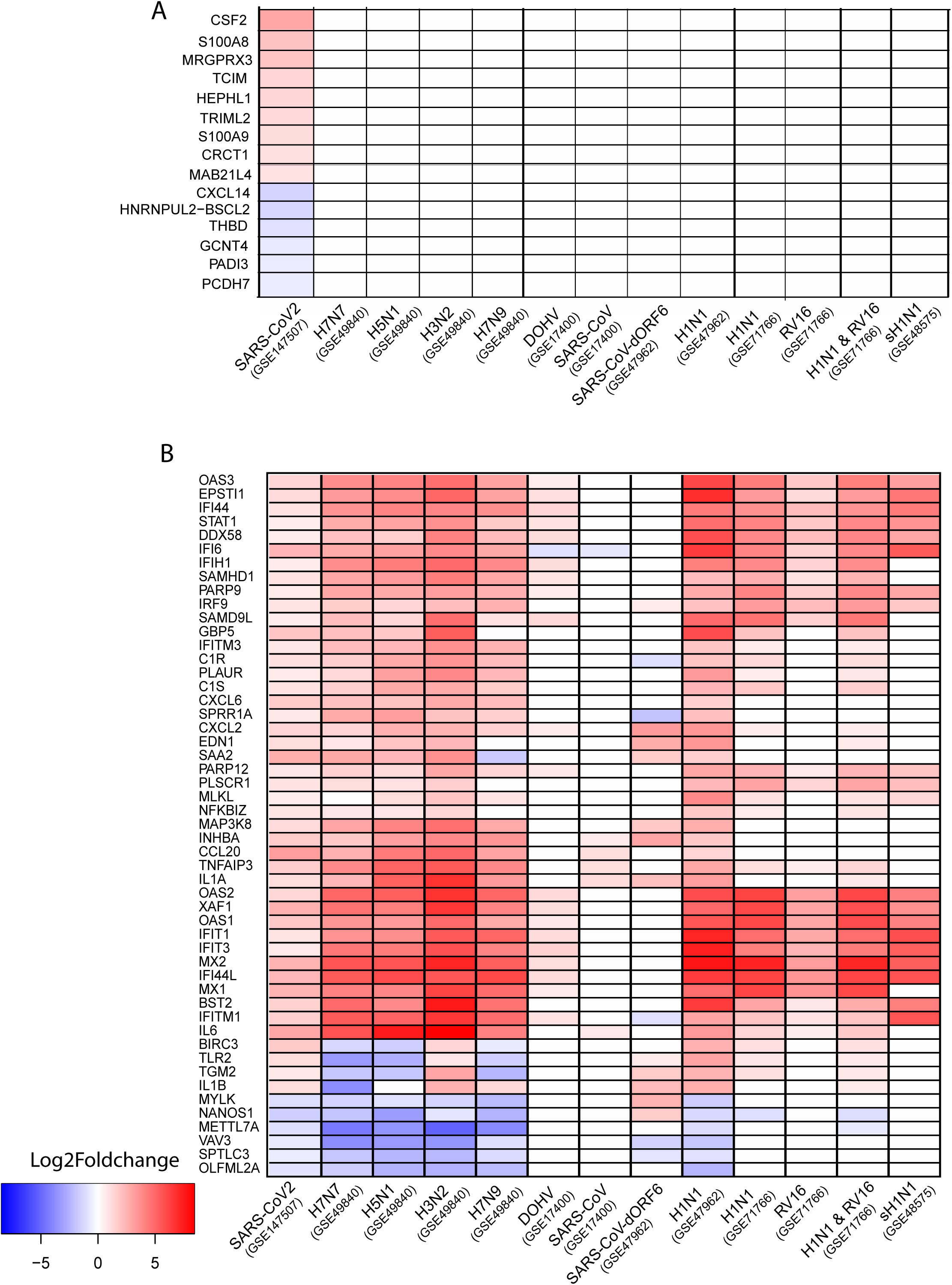
Comparative transcriptome analysis of collected gene expression data. (A) Heatmap showing 15 genes that are exclusively differentially expressed by SARS-CoV-2 infection. (B) Heatmap showing top 50 genes whose expression is altered on infection of at least two respiratory viruses including SARS-CoV-2.

Genes involved in “type1 interferon signaling pathway” and “apoptosis process” were commonly altered by SARS-CoV-2 and other viral infections. These included *OAS1, OAS2, OAS3, MX1, MX2, SAMHD1, XAF1, BST2, IFI6, IFIT1, IFIT3, IFITM1, IFITM3, IRF9, STAT1, TNFAIP3, BIRC3, IL1A, IL1B, MAP3K8, PLSCR1*, and *TLR2* [**Figure 2B**].

### Protein-protein interaction analysis of altered genes on SARS-CoV-2 infection revealed hub genes

In NHBECs, all 240 genes differentially expressed by SARS-CoV-2 infection were queried in the STRING database to identify known interactions among them. The database provided a PPI network of 1080 interactions (edges) among 168 genes (nodes) [**Figure 3**]. The PPI network was downloaded as a simple interaction format (SIF) file, visualized with Cytoscape, and analyzed with Cytohubba plugin to identify hub genes. The top 50 genes were obtained based on three network parameters: degree, closeness, and betweenness separately. The 24 genes featured in all three lists were considered as hub genes [**Table 2**]. The hub genes were *IL6, CXCL8, IL1B, STAT1, MMP9, TLR2, CXCL1, ICAM1, CSF2, IFIH1, NFKBIA, DDX58, MX1, MMP1, PI3, SAA1, BST2, LCN2, EDN1, STAT5A, C3, SOD2, LIF*, and *HBEGF*. To understand the exclusivity of these hub genes with COVID19 infection, we analyzed their expression after other viral infections. CSF2 was the only gene to remain unaltered on exposure to other viral infections; most of the COVID19 hub genes were affected by infection of human or avian influenza viruses such as *H7N7, H1N1, H7N9, H3N2*, and *H5N1* [**Figure 4**].

**Table 2:** Hub genes involved in COVID19 infection of normal human bronchial epithelial cells.

| *Hub nodes* | Degree | Closeness | Betweenness | AbsFC | adj.P-value | DrugBank |
| --- | --- | --- | --- | --- | --- | --- |
| *IL6* | 75 | 114.12 | 4521.40 | 7.60 | 6.59E-20 | Yes |
| *CXCL8* | 60 | 105.95 | 1849.39 | 5.29 | 2.64E-112 |  |
| *IL1B* | 56 | 103.70 | 2057.31 | 2.12 | 9.62E-25 | Yes |
| *STAT1* | 54 | 101.70 | 2673.02 | 1.50 | 6.69E-07 |  |
| *MMP9* | 53 | 102.83 | 3871.50 | 5.50 | 4.86E-23 | Yes |
| *TLR2* | 50 | 100.20 | 1875.00 | 2.16 | 6.55E-03 | Yes |
| *CXCL1* | 42 | 94.43 | 690.65 | 2.64 | 2.27E-38 |  |
| *ICAM1* | 39 | 94.78 | 843.10 | 3.66 | 1.24E-38 | Yes |
| *CSF2* | 38 | 94.28 | 945.23 | 7.63 | 8.01E-09 |  |
| *IFIH1* | 35 | 88.45 | 481.65 | 1.56 | 1.00E-03 |  |
| *NFKBIA* | 34 | 89.18 | 645.38 | 1.89 | 9.33E-15 | Yes |
| *DDX58* | 33 | 86.95 | 387.99 | 1.50 | 8.88E-03 |  |
| *MX1* | 31 | 85.95 | 313.72 | 5.60 | 3.06E-26 |  |
| *MMP1* | 26 | 83.77 | 695.08 | 1.76 | 1.73E-04 | Yes |
| *PI3* | 24 | 87.28 | 3786.84 | 3.72 | 4.64E-07 |  |
| *SAA1* | 23 | 81.60 | 429.27 | 5.04 | 8.62E-48 |  |
| *BST2* | 23 | 77.15 | 374.75 | 2.77 | 1.82E-03 |  |
| *LCN2* | 23 | 83.95 | 972.72 | 1.69 | 1.98E-05 |  |
| *EDN1* | 22 | 83.37 | 543.24 | 2.14 | 1.19E-08 |  |
| *STAT5A* | 21 | 84.13 | 1386.96 | 2.48 | 3.40E-03 |  |
| *C3* | 18 | 78.90 | 842.15 | 2.91 | 1.06E-41 | Yes |
| *SOD2* | 18 | 80.35 | 312.40 | 2.94 | 1.54E-41 |  |
| *LIF* | 17 | 80.13 | 376.04 | 2.52 | 1.05E-26 |  |
| *HBEGF* | 17 | 78.78 | 671.66 | 2.53 | 8.02E-27 |  |

**Figure 3:**
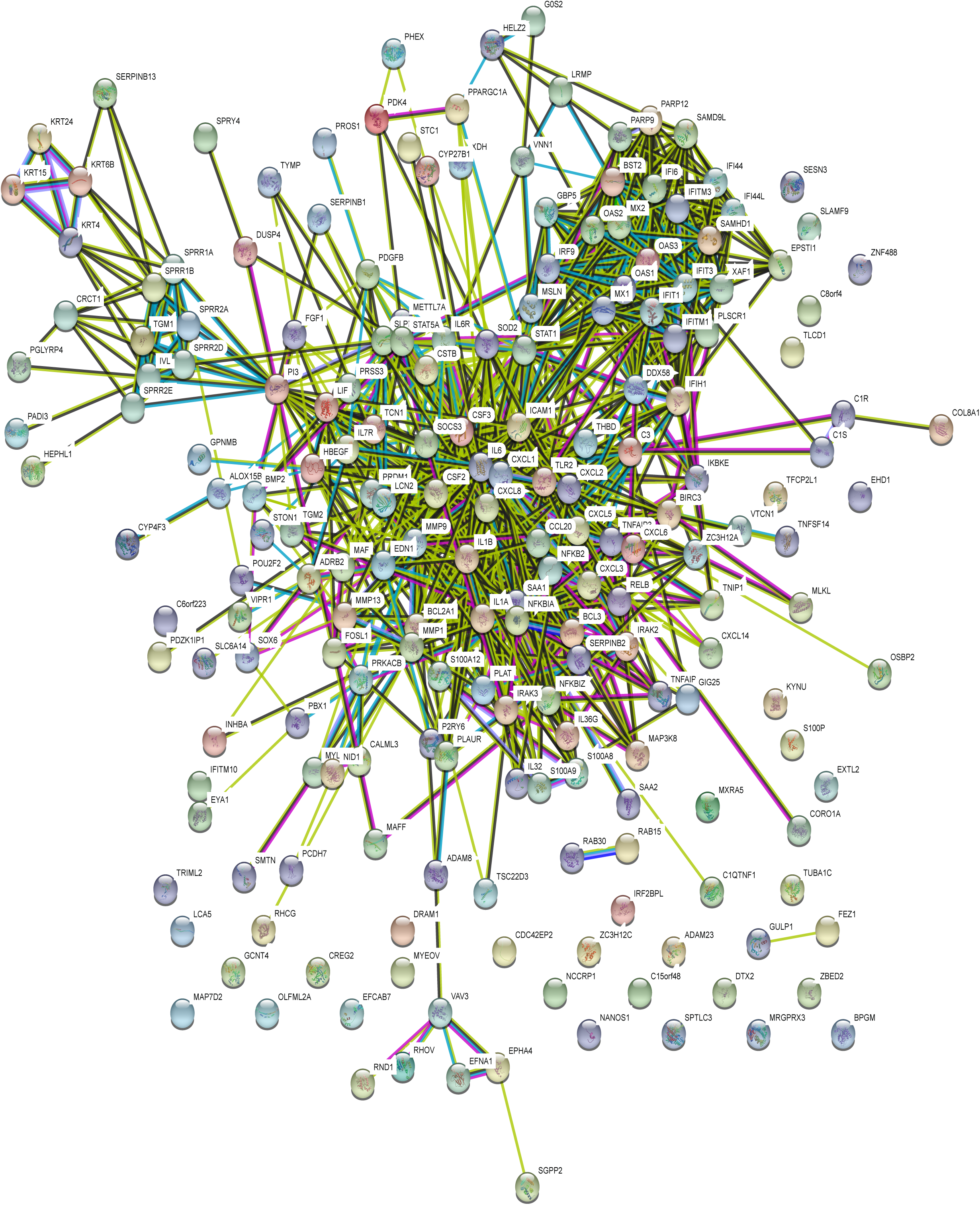
Protein-protein interaction network of genes altered by COVID-19 infection. The network with 1080 edges among 168 nodes is obtained from the STRING database. Interactions with scores of 0.4 or above were considered.

**Figure 4:**
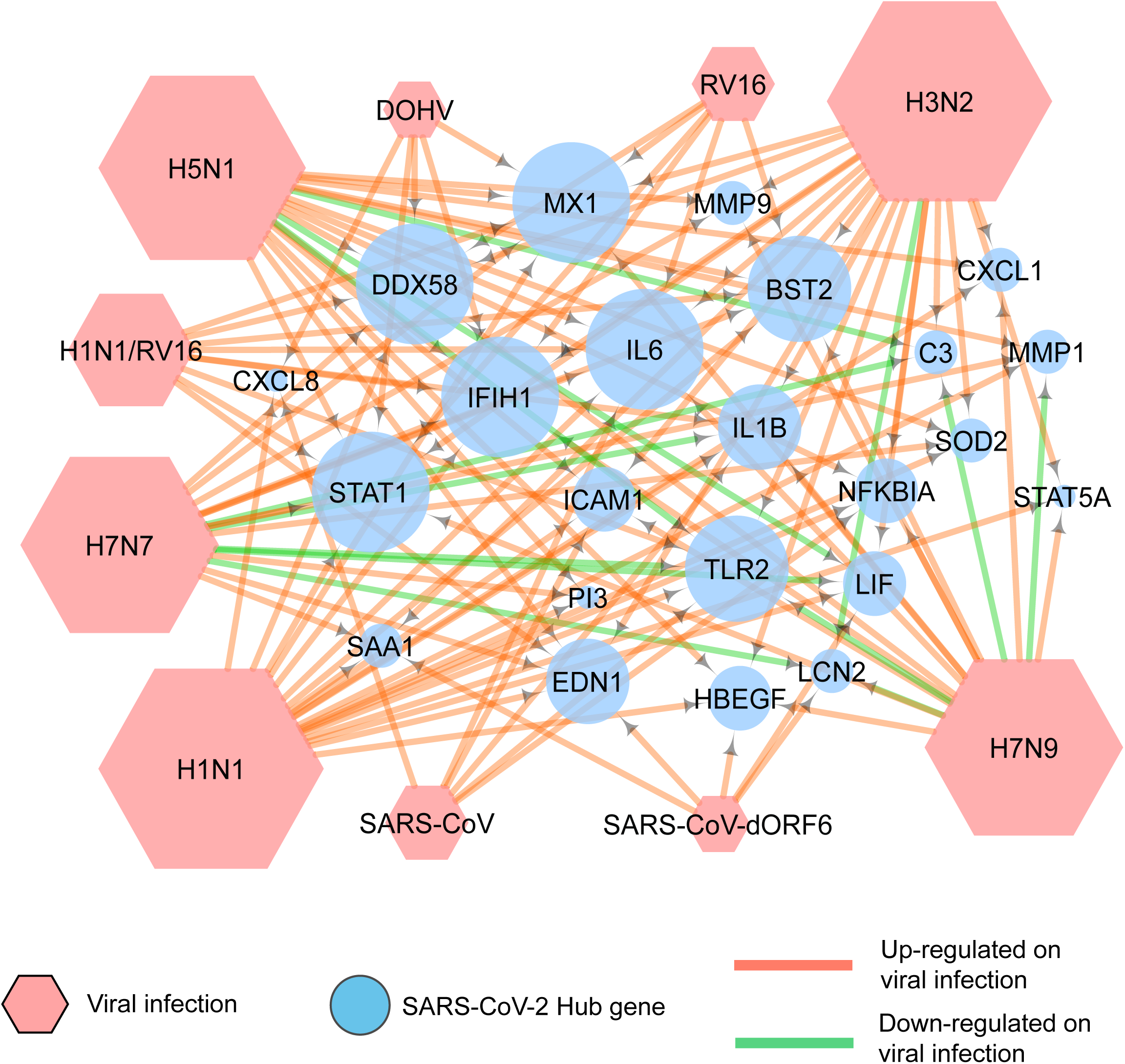
Association of COVID 19 hub genes with other respiratory viral infections. A network generated by use of Cytoscape depicts the expression of COVID-19 hub genes on infection by various respiratory viruses. The green arrows indicate down-regulation, and the red arrows show up-regulation of these genes.

## Discussion

Microarray meta-analysis and comparative transcriptome analysis have been useful bioinformatic approaches for maximum utilization of publicly available gene expression data (25, 26). Since the advent of high-throughput technologies such as microarrays and RNA-seq, researchers have performed in-depth transcriptome analysis of various biological conditions leading to multiple discoveries (27). Data from thousands of such experiments are being deposited in public repositories such as GEO and ArrayExpress. Selection and combinatorial analysis of such data can aid researchers in understanding the molecular mechanism of a disease, and in discovering biomarkers (28-30). We observed the availability of multiple microarray studies related to human respiratory viral infection and sensed the opportunity to compare the effect of SARS-CoV-2 and other respiratory viral infections on the human lung transcriptome.

The comparative transcriptome analysis led to identification of genes that are altered exclusively after SARS-CoV-2 infection. Among these genes, S100 calcium-binding proteinA9 (S100A9) and S100 calcium-binding protein A8 (S100A8) are calcium- and zinc-binding proteins that are elevated in inflammatory lung disorders (31). Colony stimulating factor 2 (CSF2) is a cytokine-coding gene associated with respiratory diseases such as pulmonary alveolar proteinosis (32). Transcriptional and immune response regulator (TCIM), a positive regulator of the Wnt/beta-catenin signaling pathway, is involved in proliferation of thyroid cancers (33, 34). Cysteine-rich C-terminal 1 (CRCT1) functions as a suppressor in esophageal cancer (35). Tripartite motif family-like 2 (TRIML2) promotes growth human oral cancers (36). Hephaestin-like 1 (HEPHL1) is a copper-binding glycoprotein with ferroxidase activity (37). MAS-related GPR family member X3 (MRGPRX3), a member of the mas-related/sensory neuron specific subfamily of G protein coupled receptors, is down-regulated in human airway epithelial cells exposed to smoke from electronic cigarettes (38). C-X-C Motif chemokine ligand 14 (CXCL14) functions as a tumor suppressor in oral cancer, lung cancer, and head and neck cancer; it also induces growth of prostate and breast cancers (39-43). Protocadherin 7 [PCDH7] is involved in cell-cell recognition and adhesion (44). Glucosaminyl (N-acetyl) transferase 4 (GCNT4), thrombomodulin (THBD), peptidyl arginine deiminase 3(PADI3), Mab-21-like 4 (MAB21L4), and HNRNPUL2−BSCL2 have no known association with lung disorders or respiratory virus infections.

The PPI analysis of genes differentially expressed by SARS-CoV-2 infection identified 24 hub genes. CSF2 was the only hub gene to show a SARS-CoV-2-exclusive gene expression pattern. Almost all other hub genes were affected by infection of other respiratory viruses. In conclusion, both PPI analysis and comparative transcriptome analysis point to a role of CSF2 in the molecular mechanism of SARS-CoV-2 infections of human lung epithelium. The current study also highlights the exclusivity of known lung inflammation disorder genes such as S100A8 and S100A9 with respect to SARS-CoV-2 infection.

Although the current bioinformatics approach makes use of available data to identify molecular targets for treatment of COVID-19, we understand that experimental validation is necessary to establish the exclusive association of CSF2, S100A8, and S100A9 with SARS-CoV-2 infection.

## Acknowledgment

This study was supported by UAB impact funds to UM and SV. We thank Dr. Donald Hill, of the UAB Comprehensive Cancer Center, for editing this manuscript.

## References

1. Carvalho A, Cezarotti Filho ML, Azevedo PCP, Silveira Filho RN, Barbosa FT, Rocha TJM, et al. Epidemiology, diagnosis, treatment, and future perspectives concerning SARS-COV-2: a review article. Rev Assoc Med Bras (1992). 2020;66(3):370–4.

2. Nicola M, Alsafi Z, Sohrabi C, Kerwan A, Al-Jabir A, Iosifidis C, et al. The socio-economic implications of the coronavirus pandemic (COVID-19): A review. Int J Surg. 2020;78:185–93.

3. Singh SK. Respiratory Viral Infections. Semin Respir Crit Care Med. 2016;37(4):485–6.

4. Afrough B, Dowall S, Hewson R. Emerging viruses and current strategies for vaccine intervention. Clin Exp Immunol. 2019;196(2):157–66.

5. Wang C, Li W, Drabek D, Okba NMA, van Haperen R, Osterhaus A, et al. A human monoclonal antibody blocking SARS-CoV-2 infection. Nat Commun. 2020;11(1):2251.

6. Blanco-Melo D, Nilsson-Payant BE, Liu WC, Uhl S, Hoagland D, Moller R, et al. Imbalanced Host Response to SARS-CoV-2 Drives Development of COVID-19. Cell. 2020;181(5):1036–45 e9.

7. Zhang N, Wang L, Deng X, Liang R, Su M, He C, et al. Recent advances in the detection of respiratory virus infection in humans. J Med Virol. 2020;92(4):408–17.

8. Parkinson H, Kapushesky M, Shojatalab M, Abeygunawardena N, Coulson R, Farne A, et al. ArrayExpress--a public database of microarray experiments and gene expression profiles. Nucleic Acids Res. 2007;35(Database issue):D747–50.

9. Barrett T, Wilhite SE, Ledoux P, Evangelista C, Kim IF, Tomashevsky M, et al. NCBI GEO: archive for functional genomics data sets--update. Nucleic Acids Res. 2013;41(Database issue):D991–5.

10. Kim D, Pertea G, Trapnell C, Pimentel H, Kelley R, Salzberg SL. TopHat2: accurate alignment of transcriptomes in the presence of insertions, deletions and gene fusions. Genome Biol. 2013;14(4):R36. 11.

11. Li H. A statistical framework for SNP calling, mutation discovery, association mapping and population genetical parameter estimation from sequencing data. Bioinformatics. 2011;27(21):2987-93. 12.

12. Anders S, Pyl PT, Huber W. HTSeq--a Python framework to work with high-throughput sequencing data. Bioinformatics. 2015;31(2):166–9.

13. Love MI, Huber W, Anders S. Moderated estimation of fold change and dispersion for RNA-seq data with DESeq2. Genome Biol. 2014;15(12):550.

14. Dennis G, Jr., Sherman BT, Hosack DA, Yang J, Gao W, Lane HC, et al. DAVID: Database for Annotation, Visualization, and Integrated Discovery. Genome Biol. 2003;4(5):P3.

15. Mitchell HD, Eisfeld AJ, Sims AC, McDermott JE, Matzke MM, Webb-Robertson BJ, et al. A network integration approach to predict conserved regulators related to pathogenicity of influenza and SARS-CoV respiratory viruses. PLoS One. 2013;8(7):e69374.

16. Kim TK, Bheda-Malge A, Lin Y, Sreekrishna K, Adams R, Robinson MK, et al. A systems approach to understanding human rhinovirus and influenza virus infection. Virology. 2015;486:146–57.

17. Yoshikawa T, Hill TE, Yoshikawa N, Popov VL, Galindo CL, Garner HR, et al. Dynamic innate immune responses of human bronchial epithelial cells to severe acute respiratory syndrome-associated coronavirus infection. PLoS One. 2010;5(1):e8729.

18. Josset L, Zeng H, Kelly SM, Tumpey TM, Katze MG. Transcriptomic characterization of the novel avian-origin influenza A (H7N9) virus: specific host response and responses intermediate between avian (H5N1 and H7N7) and human (H3N2) viruses and implications for treatment options. mBio. 2014;5(1):e01102–13.

19. Paquette SG, Banner D, Chi le TB, Leomicronn AJ, Xu L, Ran L, et al. Pandemic H1N1 influenza A directly induces a robust and acute inflammatory gene signature in primary human bronchial epithelial cells downstream of membrane fusion. Virology. 2014;448:91–103.

20. von Mering C, Jensen LJ, Snel B, Hooper SD, Krupp M, Foglierini M, et al. STRING: known and predicted protein-protein associations, integrated and transferred across organisms. Nucleic Acids Res. 2005;33(Database issue):D433–7.

21. Shannon P, Markiel A, Ozier O, Baliga NS, Wang JT, Ramage D, et al. Cytoscape: a software environment for integrated models of biomolecular interaction networks. Genome Res. 2003;13(11):2498–504.

22. Su G, Morris JH, Demchak B, Bader GD. Biological network exploration with Cytoscape 3. Curr Protoc Bioinformatics. 2014;47:8 13 1–24.

23. Chin CH, Chen SH, Wu HH, Ho CW, Ko MT, Lin CY. cytoHubba: identifying hub objects and sub- networks from complex interactome. BMC Syst Biol. 2014;8 Suppl 4:S11.

24. Wishart DS, Knox C, Guo AC, Cheng D, Shrivastava S, Tzur D, et al. DrugBank: a knowledgebase for drugs, drug actions and drug targets. Nucleic Acids Res. 2008;36(Database issue):D901–6.

25. Ramasamy A, Mondry A, Holmes CC, Altman DG. Key issues in conducting a meta-analysis of gene expression microarray datasets. PLoS Med. 2008;5(9):e184.

26. Hamid JS, Hu P, Roslin NM, Ling V, Greenwood CM, Beyene J. Data integration in genetics and genomics: methods and challenges. Hum Genomics Proteomics. 2009;2009.

27. Cahan P, Rovegno F, Mooney D, Newman JC, St Laurent G, 3rd, McCaffrey TA. Meta-analysis of microarray results: challenges, opportunities, and recommendations for standardization. Gene. 2007;401(1-2):12–8.

28. Chen JA, Yu Y, Xue C, Chen XL, Cui GY, Li J, et al. Low microRNA-139 expression associates with poor prognosis in patients with tumors: A meta-analysis. Hepatobiliary Pancreat Dis Int. 2019;18(4):321–31.

29. Sherafatian M, Abdollahpour HR, Ghaffarpasand F, Yaghmaei S, Azadegan M, Heidari M. MicroRNA Expression Profiles, Target Genes, and Pathways in Intervertebral Disk Degeneration: A Meta-Analysis of 3 Microarray Studies. World Neurosurg. 2019;126:389–97.

30. Huang W, Ran R, Shao B, Li H. Prognostic and clinicopathological value of PD-L1 expression in primary breast cancer: a meta-analysis. Breast Cancer Res Treat. 2019;178(1):17–33.

31. Gomes LH, Raftery MJ, Yan WX, Goyette JD, Thomas PS, Geczy CL. S100A8 and S100A9-oxidant scavengers in inflammation. Free Radic Biol Med. 2013;58:170–86.

32. Ito M, Nakagome K, Ohta H, Akasaka K, Uchida Y, Hashimoto A, et al. Elderly-onset hereditary pulmonary alveolar proteinosis and its cytokine profile. BMC Pulm Med. 2017;17(1):40.

33. Chua EL, Young L, Wu WM, Turtle JR, Dong Q. Cloning of TC-1 (C8orf4), a novel gene found to be overexpressed in thyroid cancer. Genomics. 2000;69(3):342–7.

34. Sunde M, McGrath KC, Young L, Matthews JM, Chua EL, Mackay JP, et al. TC-1 is a novel tumorigenic and natively disordered protein associated with thyroid cancer. Cancer Res. 2004;64(8):2766–73.

35. Wu N, Song Y, Pang L, Chen Z. CRCT1 regulated by microRNA-520 g inhibits proliferation and induces apoptosis in esophageal squamous cell cancer. Tumour Biol. 2016;37(6):8271–9.

36. Hayashi F, Kasamatsu A, Endo-Sakamoto Y, Eizuka K, Hiroshima K, Kita A, et al. Increased expression of tripartite motif (TRIM) like 2 promotes tumoral growth in human oral cancer. Biochem Biophys Res Commun. 2019;508(4):1133–8.

37. Sharma P, Reichert M, Lu Y, Markello TC, Adams DR, Steinbach PJ, et al. Biallelic HEPHL1 variants impair ferroxidase activity and cause an abnormal hair phenotype. PLoS Genet. 2019;15(5):e1008143. 38.

38. Solleti SK, Bhattacharya S, Ahmad A, Wang Q, Mereness J, Rangasamy T, et al. MicroRNA expression profiling defines the impact of electronic cigarettes on human airway epithelial cells. Sci Rep. 2017;7(1):1081.

39. Ozawa S, Kato Y, Komori R, Maehata Y, Kubota E, Hata R. BRAK/CXCL14 expression suppresses tumor growth in vivo in human oral carcinoma cells. Biochem Biophys Res Commun. 2006;348(2):406–12.

40. Augsten M, Hagglof C, Olsson E, Stolz C, Tsagozis P, Levchenko T, et al. CXCL14 is an autocrine growth factor for fibroblasts and acts as a multi-modal stimulator of prostate tumor growth. Proc Natl Acad Sci U S A. 2009;106(9):3414–9.

41. Ozawa S, Kato Y, Ito S, Komori R, Shiiki N, Tsukinoki K, et al. Restoration of BRAK / CXCL14 gene expression by gefitinib is associated with antitumor efficacy of the drug in head and neck squamous cell carcinoma. Cancer Sci. 2009;100(11):2202–9.

42. Tessema M, Klinge DM, Yingling CM, Do K, Van Neste L, Belinsky SA. Re-expression of CXCL14, a common target for epigenetic silencing in lung cancer, induces tumor necrosis. Oncogene. 2010;29(37):5159–70.

43. Augsten M, Sjoberg E, Frings O, Vorrink SU, Frijhoff J, Olsson E, et al. Cancer-associated fibroblasts expressing CXCL14 rely upon NOS1-derived nitric oxide signaling for their tumor-supporting properties. Cancer Res. 2014;74(11):2999–3010.

44. Nakamura H, Nakashima T, Hayashi M, Izawa N, Yasui T, Aburatani H, et al. Global epigenomic analysis indicates protocadherin-7 activates osteoclastogenesis by promoting cell-cell fusion. Biochem Biophys Res Commun. 2014;455(3-4):305–11.

